# Exploring different sequence representations and classification methods for the prediction of nucleosome positioning

**DOI:** 10.1101/482612

**Authors:** Nikos Kostagiolas, Nikiforos Pittaras, Christoforos Nikolaou, George Giannakopoulos

**Affiliations:** Information Management Systems Institute, “Athena” Research & Innovation Center, Athens, 15125, Greece; Institute of Informatics and Telecommunications, NCSR “Demokritos”, Athens, 15341, Greece; Department of Biology, University of Crete, Heraklion, 70013, Greece

## Abstract

**Motivation:** Nucleosomes form the first level of DNA compaction and thus bear a critical role in the overall genome organization. At the same time, they modulate chromatin accessibility and, through a dynamic equilibrium with other DNA-binding proteins, may shape gene expression. A number of large-scale nucleosome positioning maps, obtained for various genomes, has compelled the importance of nucleosomes in the regulation of gene expression and has shown constraints in the relative positions of nucleosomes to be much stronger around regulatory elements (i.e. promoters, splice junctions and enhancers). At the same time, the great majority of nucleosome positions appears to be rather flexible. Various computational methods have in the past been used in order to capture the sequence determinants of nucleosome positioning but, as the extent to which DNA sequence preferences may guide nucleosome occupancy largely varies, this has proved to be rather difficult. In order to focus on highly specific sequence attributes, in this work we have analyzed two well-defined sets of nucleosome-occupied sites (NOS) and nucleosome-free-regions (NFR) from the genome of *S. cerevisiae*, with the use of textual representations.

**Results:** We employed 3 different genomic sequence representations (Hidden Markov Models, Bag-of-Words and N-gram Graphs) combined with a number of machine learning algorithms on the task of classifying genomic sequences as nucleosome-free (NFR) or nucleosome-occupied NOS (to be further amended based on updated results). We found that different approaches that involve the usage of different representations or algorithms can be more or less effective at predicting nucleosome positioning based on the textual data of the underlying genomic sequence. More interestingly, we show that N-gram Graphs, a sequence representation that takes into account both k-mer occurrences and relative positioning at various lengths scales is outperforming other methodologies and may thus be a choice of preference for the analysis of DNA sequences with subtle constraints.

## 1 Introduction

### 1.1 Background

Nucleosomal particles are formed through the binding of a histone octamer around a 147-nucleotide stretch of eukaryotic DNA. They constitute the basic component of chromatin packaging and cover the greatest part (between 75 and 90 per cent) of all eukaryotic genomes. In this way, they modulate chromatin accessibility and compete with other DNA-binding factors in shaping the regulatory potential of many genes. Even though mapping nucleosome positions genome-wide is nowadays possible through various high-throughput approaches [1, 2, 3, 4], it is well established that the large majority of nucleosome positions in a given cell type and for given conditions are highly flexible, occupying space in a non-reproducible manner [5, 6]. On the other hand, a subset of the total nucleosomes tend to be positioned with high specificity around genomic regions of regulatory role such as around transcription start and end sites, promoters [1, 7], splice junctions [8] and enhancers [9].

This well-documented link between nucleosomes and transcriptional regulation has driven efforts to define the sequence determinants that underly nucleosome positioning. Initial computational approaches on this problem were based on probabilistic models of position-specific dinucleotide composition of in vivo [10] or in vitro [11] formed nucleosomes. Even though position-specific constraints do exist in nucleosome sequences they are rather weak and so methods using more generalized sequence content analysis focused at the discovery of periodical patterns with the use of wavelets [12], duration Hidden Markov Models [13] or by simply applying periodicity analysis on structural data [14]. Other methods make use of biophysical [15], statistical mechanics models [16] or structural constraints [8] in order to quantify a given sequence’s affinity to form nucleosomes. A general characteristic of all such attempts was that they overall performed poorly when tested genome-wide [17]. Even though various computational methods had been utilized, varying from likelihood models [11] [10], to supervised learning strategies [18, 19], the entirety of the results would converge to the same conclusion that there is little consistency in nucleosome positioning across the eukaryotic cellular population. This is due to the underlying statistical positioning, a concept initially put forward on the basis of theoretical calculations [20], but which has nowadays been supported through modeling of nucleotide content [21], comparative analyses of public data [22] or experimentally, through the coupling of nucleosome positioning with very deep DNA sequencing [6].

### 1.2 Motivation

Besides providing valuable insight in the way chromatin affects basic cellular processes, the definition of the sequence determinant driving nucleosome positioning, may have broader implications as recent works [23] have described experimental approaches for the engineering of genomic sequences with particular structural properties. In [22], we showed constitutively (i.e. highly reproducible) nucleosome positioning and nucleosome-free regions to bear specific properties in terms of DNA structure and sequence conservation. In the same work, we provided support of a model according to which, NFRs may act as the organizing principle for statistical positioning. We found constitutive yeast NFRs, defined through a combination of three individual nucleosomal maps [1, 24, 25] to be preferentially positioned close to housekeeping genes and with increased sequence conservation. We were also able to define well-positioned nucleosomes in the vicinity of (or immediatelly flanking) these NFRs and, more importantly, showed these “constitutive” nucleosomes to carry particular constraints revealed at the structural level of conserved DNA curvature patterns. In this work, we follow up on these two sets of sequences in order to discover particular sequence characteristics that can be attributed to its tendency to form or exclude nucleosomes.

Research has so far resorted to the established methods of Hidden Markov Models (HMMs) [13] and Support Vector Machines [26], without being able to capture nucleosome-forming sequence constraints besides fundamental compositional tendencies (e.g. GC content). It would be thus, interesting to examine to what extent the adoption of a broader selection of Machine Learning algorithms and representations would be able to produce better results towards predicting nucleosome positions while at the same time revealing certain patterns in the textual data of the underlying genomic sequence that inhibit or promote nucleosome positioning. This work addresses this question by introducing the application of a variety of Machine Learning algorithms (*Decision Trees, Naive Bayes, k-Nearest Neighbors, Support Vector Machines*) and representations (*Hidden Markov Models, N-gram Graphs, Bag-of-Words*), thus supporting varying use of contextual information. This approach allows for a more complete evaluation spectrum of several possible algorithm-representation pairs to be established through a unified set of experiments and evaluations.

## 2 Methods

### 2.1 Problem Definition

The goal of our work is to define the principles that regulate the formation or absence of nucleosomes within the underlying genomic sequence while also providing a comprehensive overview of different feature extraction pipelines with the introduction of the appropriate representation spaces, different algorithms and metrics. In order to achieve this, we opt to solve a single-label binary classification problem, by predicting whether the textual elements of a genomic sequence favor the formation of nucleosome occupied site or not. Therefore, the genomic sequences in our dataset belong either to the NFR class (Nucleosome-Free Region) or to the NOS class (Nucleosome Occupancy Site).

We stress that we will represent the textual elements in a variety of manners. Thus, we expect that different types of representation will highlight different textual traits to guide the classification process. As a consequence, the classification itself is used as the means to: understand the connection between different representations and learning algorithms and the classification outcomes; identify promising and robust combinations that can be repeatedly used in future analyses.

### 2.2 Proposed Method

In this section we outline the proposed method for nucleosome positioning prediction, consisting of two parts: i) feature representation, where the input sequence is transformed in a computationally convenient format; ii) Classification, where the representations from (i) are used to build a classification model, classifying the sequence as an NFR or NOS.

In the problems we face, we consider that we have a number of labeled, training sequences and we need to predict the class in a number of held-out, test sequences. We want to understand whether different representation models offer varying predictive power in the given problem, as has been noted in other classification problems [27].

We have followed two approaches for the representation, as follows:

- *Instance-based:* In this approach each sequence is defined by its content properties (e.g. frequency of k-mers), as is commonly applied. The resulting feature space is a vector space, following the Vector Space Model (VSM) paradigm [28].
- *Class-representatives-based:* In this approach we first create one representative per class, trying to encapsulate all the information related to all the instances of the class (e.g. centroid of class instances). Then, all instances are described by how similar they are to these representatives, via one or more similarity measures (e.g. cosine similarity, euclidean distance). In this case, the resulting feature space is still a vector space, but the dimensions represent similarities to representatives and not self-sufficient content information. This view allows us to employ models, such as Hidden Markov Models and N-gram Graphs — which encode non-vectorial, sequential or neighborhood patterns — but still get a vector space, usable by most established classification methods.

In the following paragraphs we elaborate on the exact representations used.

### 2.2.1 Bag-of-words

The Bag-of-words (BoW) model [29, 30] is a VSM able to represent sequences as the multiset of selected tokens/subsequences. We term this multiset the “bag”. Despite keeping tokens which occur multiple times in a document, the BoW model ignores any form of structure present in the sequence, including token order. A typical BoW encoding produces a sparse vector where each coordinate corresponds to a subsequence, and its value holds the number of times that subsequence appears in the input text. In our case, we create a bag-of-nucleotides representation for a given genomic sequence by counting the occurrences of each nucleotide in the input sequence.

In order to embed information about k-mers, i.e. go beyond individual nucleotides, we consider k-mers as subsequences of length *k* ∈ ℕ*\**. We adopt the widely used values of *k* = 3 (in Natural Language Processing) and *k* = 2 (in genomic studies similar to this one) with a rolling window step of 1 (resulting in overlapping 3-mers or 2-mers), parameters that present a good trade-off between model complexity and computational cost.

We apply the BoW model in two variations, as follows.

1. **Baseline-BoW:** In this approach, we use the raw frequencies of each 3-mer in the input sequence, with two normalizations: First, we compute the relative frequency by dividing by the number of possible k-mers in the genomic code. This normalization, apart from scaling and correcting frequency imbalance, produces unit-length vectors, which is often advantageous in machine learning classification. Secondly, we apply a correction for background nucleotide composition, dividing the relative k-mer frequency in the sequence, by the product of the relative frequencies of the 1-mers that compose that k-mer; This is due to the DNA composition at mono-nucleotide level being known to be highly variable along the higher eukaryotic genome. For example, if our input language *L* consists of all possible 3-mers producible over the genomic alphabet {*A, C, G, T*}, then |*L*| = 4^3^ = 64 and given a genomic sequence instance *g*, we end up with the following feature vectors :

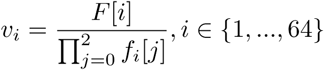

where *v*_*i*_ is the *i*-th coordinate, *F* [*i*] is the relative frequency of the *i*-th 3-mer in the the instance sequence, *f*_*i*_[*j*] is the relative frequency of the *j*-th 1-mer of the *i*-th 3-mer, with respect to the sequence length. It is apparent that this type of feature extraction results in multidimensional feature vectors in ℝ^64^, where each dimension is mapped to a 3-mer. This approach is clearly an instance-based appraoch.
2. **Centroid-BoW:** This approach is class-representative-based. Thus, we apply a preprocessing phase to our model, using the majority of the training data to extract centroid vectors for our two classes. To position any instance (whether training or test) in the similarity-based space of the Centroid BoW, we compute the cosine similarity of the instance representation and each class representative vector. This results in a final feature vector *v* ∈ ℝ^2^ for Centroid-BoW, with the first (second) coordinate in *v* representing the similarity of the instance with the NFR (NOS) class. To demonstrate the process, consider the following eight genomic sequences, four of them belonging to the NFR class and four of them to the NOS class, and the extracted instance bags:

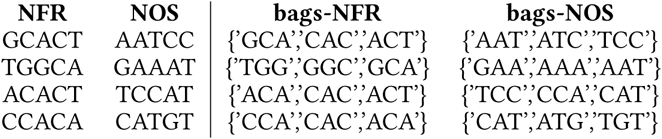 The NFR and NOS class vectors are then formed, by merging the 3-mers that belong to the same class and computing the relative frequencies of the distinct 3-mers:

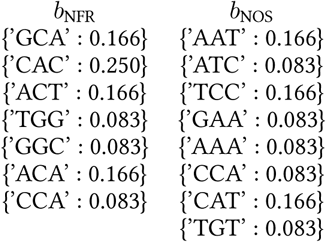 We note that in all the cases where we describe a vector, we omit the dimensions that have zero values, for brevity (i.e. we use a sparse representation). Now let us position two example instances in the vector space: *q*_1_ = **CGCAAATTATTG** and *q*_2_ = **AGCAAGACAGTT**. Their corresponding relative-frequency bags of 3-mers are the following:

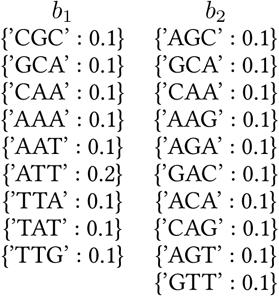 The cosine similarity between *q*_1_ and the NFR representation is:

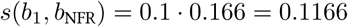

while the similarity to the NOS bag is :

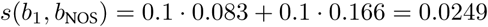 Following the same process we compute the cosine similarities between *q*_2_ and the class vectors, resulting in the following feature vectors for *q*_1_ and *q*_2_:

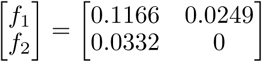 As can be seen, this approach differs from the baseline variant, because the features produced are based on the similarity of the bag of each instance with the two class centroid bags. The centroid-based approach also introduces significant *dimensionality reduction* to our problem by transforming the feature space from a 64-dimensional model to a 2-dimensional one.

### 2.2.2 Hidden Markov Models - HMMs

Hidden Markov Models (HMMs) [31] are a common baseline statistical model used in sequential or temporal environments. An HMM is a statistical Markov model in which the system being modeled is assumed to be a Markov process with unobserved (hidden) states. However, despite not having knowledge about the state sequence that the model transitions through, that specific information is provided through a probability function which describes the evolution of the system.

Similar to the centroid-BoW model, we use HMMs to describe the whole set of instances per class (NFR and NOS). Thus, we create an HMM for the NFR training instances and an HMM for the NOS ones. Then, we describe each instance (whether training or testing) by measuring the likelihood of the instance, given the corresponding model. This likelihood essentially acts as a similarity measure between an instance and the corresponding HMM model. Hence, given a nucleotide sequence *q* and two HMMs *H*_NFR_ and *H*_NOS_ belonging to the NFR and NOS classes respectively, the feature vector representation *v* of *q* will consist of these two probability values:

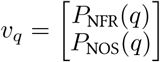

where *P*_*C*_ (*q*) is the output probability that the sequence *q* is generated by the HMM of class *C*, where *C* ∈ {NFR, NOS}. We illustrate the HMM-based feature extraction process with the following example :

#### Example 2.1

*Consider the two genomic sequences q*_1_ = *CGCAAATTATTG and q*_2_ = *AGCAAGACAGTT, Having trained an HMM model for each class, we proceed to compute the conditional probabilities of each test sequence belonging to each and class by evaluating the HMM models, producing the following representations:*

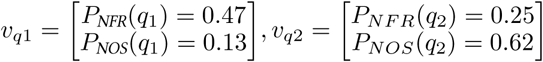

*Hence, the feature vector for each of the two sequences corresponds to a row in the following matrix:*

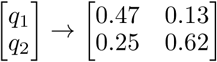

In addition, we also include a variant of feature extraction for the same HMM model, entitled **Normalized HMM**, following the suggestions in *Antonakaki et. al* [32]. In the normalized case, the probability values are passed through a logarithm function and are subsequently normalized by division with the nucleotide sequence length, |*q*|. Specifically, our feature vector *v* will now have the following form:

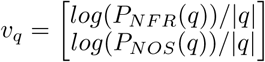

The reasons behind this approach are linked to the fact that the HMM output probabilities are a result of a product of marginal probabilities, computed in the Markov chain of the model. As a result, the output value is inversely proposal to the nucleotide sequence length. In other words, as the length of the input sequence increases, its probability decreases in an analogous manner resulting in skewed probability values across sequences with different lengths. To counter this, we first normalize the using the logarithm function to achieve numerical stability and proceed to normalize it using the division by the sequence length, as described above.

### 2.2.3 N-gram Graphs

Another useful model which can be used to represent text data is the n-gram graph. In text-related studies, an n-gram is a set which consists of *n* consequtive characters or words. The n-gram graph model originating from Natural Language Processing [33], has been successfully applied to a variety of settings, including text summarization and classification [34] but also genomic sequence analysis [27].

An *n-gram graph* is a graph *G* = {*V ^G^, E^G^*} the vertices of which, *v*^*G*^ ∈ *V ^G^*, are n-grams, and the edges *e*^*G*^ ∈ *E*^*G*^ connecting them represent adjacency frequency between the corresponding n-grams. A formal definition is given by Giannakopoulos *et al.* [33]:

#### Definition 2.1

“*If S* = {*S*_1_, *S*_2_, *…*}, *S*_*k*_ ≠ *S*_l_, ∀*k* ≠ *l, k, l* ∈ ℕ *is the set of distinct n-grams extracted from a text T^l^, and S_i_ is the i-th extracted n-gram, then G is a graph, where there is a bijection (one-to-one and onto) function f* : *S → V.”*

The edges *E*, are labeled with weights *c*_*ij*_ where *c*_*ij*_ corresponds to the number of times an adjacency of a specific pair of n-grams *S*_*i*_, *S*_*j*_ grams occurs within a specified window *D*_*win*_ of each other. The window size is measured, in our case, as the number of nucleotides from the beginning to the end of the window. In our study, given a sequence, we construct the n-gram graph applying the symmetric approach, as described in Giannakopoulos et al.[33].

#### Definition 2.2

***The symmetric approach*** *-“A window of length n runs over the summary text, centered at the beginning of N*_0_. *If the n-gram we are interested in is located at position p*_0_, *then the window will span from* 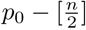 *to* 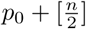 *taking into account both preceding and following characters or words. Each neighbourhood indicative edge is weighted based on the number of window co-occurrences of the neighbours, as previously indicated, within the text.”*

In our, genomic setting the n-gram graphs essentially provide an overall picture of local (within-neighborhood) phenomena of *k*-mer co-occurrence. This representation contains more sequence-related information thatn the BoW model, but is less restrictive than the HMMs in terms of the exact order of *k*-mers in the sequence. Thus, we expect n-gram graphs to combine sequential information with a liberal view of co-occurrence to describe mid-range dependencies and phenomena.

In the n-gram graph representation, we can represent sets of sequences through two steps: we first represent each sequence as a graph; we then apply the merging/update operator [35] multiple times to extract a “centroid-like” graph. Then, to position any instance in a common vector space: we construct an n-gram graph for the sequence; we compare the graph with the representative class graphs. The similarities can then be assigned to the dimensions of our feature vector.

In this study we use three forms of similarity metrics between two n-grams *G*^*i*^, *G*^*j*^: the Value Similarity (or *V S*) which measures the amount of edges which are contained both in graph *G*^*i*^ and in graph *G*^*j*^, the Size Similarity (or *SS*) which compares the size between *G*^*i*^, *G*^*j*^ and the Containment Similarity (or *CS*) which measures the part of *G*^*i*^ contained by *G*^*j*^. Here we will provide the formal definitions fo each metric [35]:

#### Definition 2.3

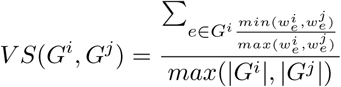

#### Definition 2.4

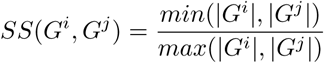

#### Definition 2.5

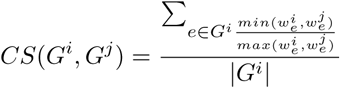

where *G*^*i*^, *G*^*j*^ are the graphs we compare; 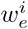 is the weight of edge *e* in graph *G*^*i*^, |*G*| is the number of edges in graph *G*.

Hence, our vector *v*(*i*) of the sequence *i* will be of the form:

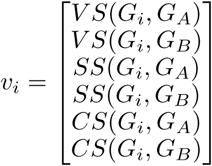

where *G*_*i*_ is the n-gram graph extracted from the genomic sequence instance *i*, while *G*_*A*_, *G*_*B*_ are class-representatives for class *A, B* correspondinly.

## 3 Results

In the following paragraphs we elaborate on the experimental setup and the evaluation of the studied methods.

### 3.1 Experimental Setup

The dataset we will use for our study consists of the *S. cerevisiae* genome and is a similar dataset to that used in [22]. Due to it being a commonly used dataset in previous studies, we can easily compare our results with that of previous experiments. However, it is important to note that the aforementioned dataset exhibits class imbalance, as it includes only 1033 instances (comprising roughly 28% of the dataset) for the NFR class opposed to the 2660 instances (constituting the rest 72% of the dataset) for the NOS class. For conducting our experiments we used 10-fold cross-validation. The final results reported in Section 3.2 are the averages of the metrics obtained for every different training-validation split.

For handling the Hidden Markov Model part of the implementation, Jahmm^3^, a Java library implementing the various algorithms related to HMMs was used. For training our HMM model, we used the Baum-Welch algorithm [36], which was implemented in this library.

In addition, for the n-gram graph part of the implementation, the JINSECT^4^ toolkit was used. JINSECT is a Java-based toolkit and library that supports and demonstrates the use of the n-gram graphs representation and the related similarity algorithms.

Finally, we used Weka^5^ [37] a collection of machine learning algorithms for data mining tasks, in order to utilize these algorithms for our classification tasks. The classifiers used where Decision Trees, k-Nearest Neighbors, Naive Bayes and SVMs.

### 3.2 Evaluation

We hereby present our experimental results. We used the measures Recall, Precision and F-measure [38] to evaluate the performance of our experiments. For brevity, we only elaborate on F-measure results — Figure 1 and Tables 1, 4 — in this paper, since it takes equally into account precision and recall. Later, we also provide some additional information of the relative ranking of methods with respect to precision and recall, in Tables 3, 7, 8.

**Table 1:** Mean F-measure per Algorithm/Representation Pair (higher is better). Best measurements per representation appear underlined while the best measure overall appears in bold.

| Representation | Classification Algorithm | Mean F-measure |
| --- | --- | --- |
| HMM | Naive Bayes | 0.764 |
|  | Decision Trees | <u>0.839</u> |
|  | kNN | 0.809 |
|  | SVM | <u>0.839</u> |
| Normalized HMM | Naive Bayes | 0.888 |
|  | Decision Trees | <u>0.890</u> |
|  | kNN | 0.810 |
|  | SVM | 0.868 |
| Centroid Bag-of-words | Naive Bayes | 0.947 |
|  | Decision Trees | <u>0.963</u> |
|  | kNN | 0.961 |
|  | SVM | 0.945 |
| Normalized Bag-of-words | Naive Bayes | 0.951 |
|  | Decision Trees | 0.982 |
|  | kNN | 0.964 |
|  | SVM | **0.984** |
| N-gram graphs | Naive Bayes | 0.969 |
|  | Decision Trees | 0.972 |
|  | kNN | <u>0.974</u> |
|  | SVM | 0.971 |

**Table 2:** Mean PRC Area scores for worse-than-chance Representation / Classification algorithm pairs. Measurements for each class are shown (including their weighted average)

| Representation | Classifier | Class | Mean PRC Area |
| --- | --- | --- | --- |
| HMM | k-Nearest Neighbors | NFR | 0.398 |
|  |  | NOS | 0.786 |
|  |  | Weighted avg. | 0.678 |
|  | Naive Bayes | NFR | 0.441 |
|  |  | NOS | 0.894 |
|  |  | Weighted avg. | 0.768 |
| Normalized HMM | k-Nearest Neighbors | NFR | 0.406 |
|  |  | NOS | 0.797 |
|  |  | Weighted avg. | 0.688 |

**Table 3:** Comparison between bigram and trigram Bag-of-words models based on the mean Recall, Pre-cision and F-measure metrics (higher is better). Best measurements per classifier-metric pair appear underlined while the best measure overall appears in bold.

| Metric | Classifier | 2-gram | 3-gram |
| --- | --- | --- | --- |
| Recall | Naive Bayes | **1.000** | **1.000** |
|  | Decision Trees | <u>0.983</u> | 0.947 |
|  | kNN | <u>0.982</u> | 0.970 |
|  | SVM | **1.000** | **1.000** |
| Precision | Naive Bayes | 0.945 | 0.945 |
|  | Decision Trees | <u>0.963</u> | 0.944 |
|  | kNN | <u>0.973</u> | 0.971 |
|  | SVM | 0.978 | **0.981** |
| F-measure | Naive Bayes | 0.951 | 0.951 |
|  | Decision Tree | 0.961 | <u>0.982</u> |
|  | kNN | <u>0.971</u> | 0.964 |
|  | SVM | 0.981 | **0.984** |

**Table 4:** ANOVA results for the F-measure metric.

|  | Df | Sum Sq | Mean Sq | F value | Pr(>F) |
| --- | --- | --- | --- | --- | --- |
| Representation | 4 | 1.8248 | 0.4562 | 3836.988 | < 2e-16 |
| Classifier | 3 | 0.0053 | 0.0018 | 14.939 | 1.28e-08 |
| Fold | 1 | 0.0003 | 0.0003 | 2.396 | 0.124 |
| Representation:Classifier | 12 | 0.0443 | 0.0037 | 31.063 | < 2e-16 |
| Representation:Fold | 4 | 0.0004 | 0.0001 | 0.843 | 0.500 |
| Classifier:Fold | 3 | 0.0002 | 0.0001 | 0.441 | 0.724 |
| Representation:Classifier:Fold | 12 | 0.0007 | 0.0001 | 0.479 | 0.925 |
| Residuals | 160 | 0.0190 | 0.0001 |  |  |

**Figure 1:**
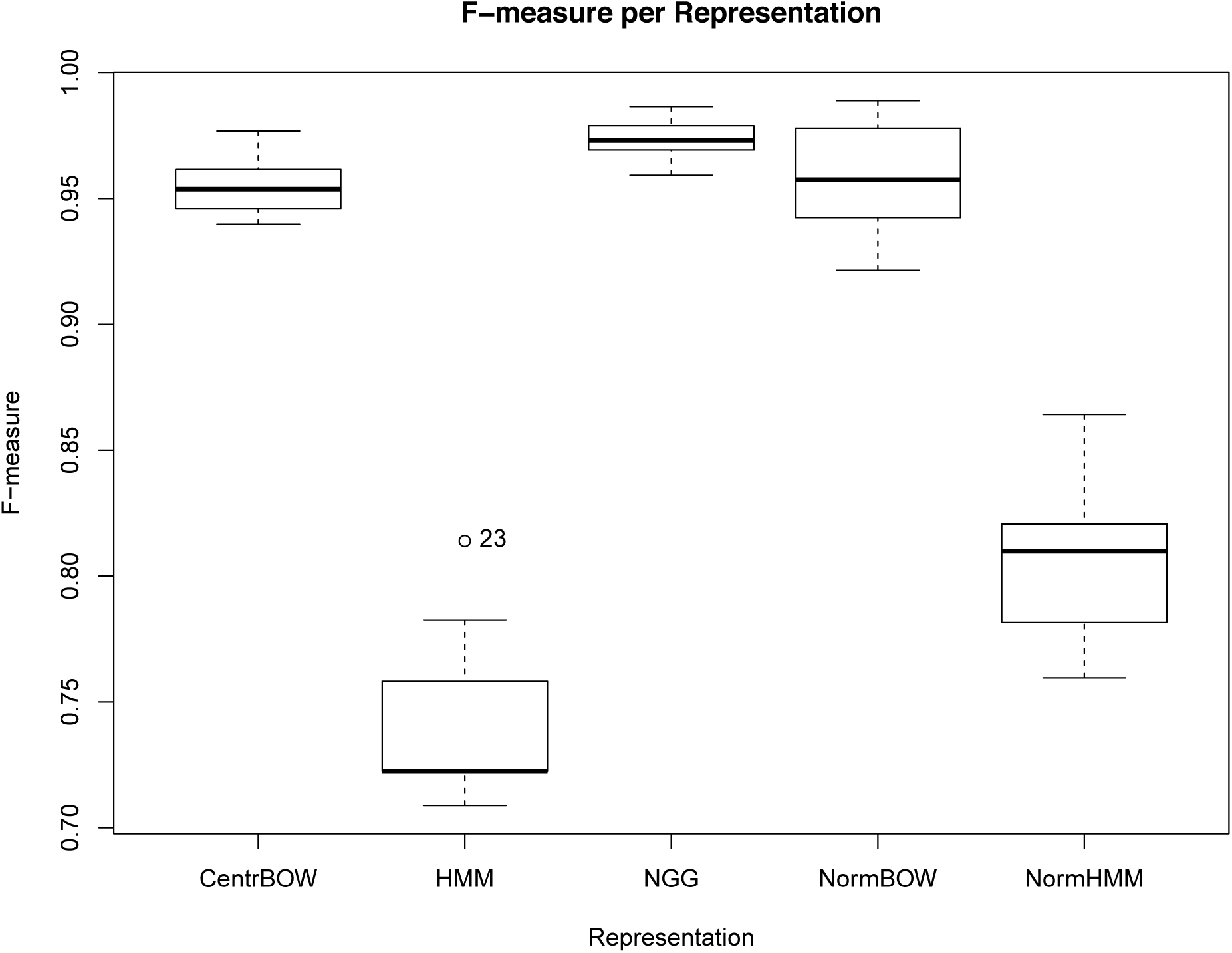
Boxplots on the F-measure per Representation. *NOTE: The y-axis holds values between 0.70 and 1.00 for better visualization of the differences.*

The most successful models in terms of F-measure are the Normalized Bag-of-words and N-gram graphs models, the mean F-measure of which resides at 0.984 and 0.974 respectively. HMMs tend to be the least successful ones featuring a mean F-measure of 0.812. Normalized HMMs appear to be more effective than plain ones with a mean F-measure at 0.864, while the Centroids Bag-of-words model featuring dimensionality reduction tends to be less successful than the Normalized Bag-of-words model by a mean margin of 0.004.

A thing to note is that, given the number of positive and negative examples, a fully random classifier would have a 72% chance of being correct, thus leading to a maximum F-measure score of 84%. However, due to some Representation-Classification algorithm pairs performing below this score (namely the combinations of HMMs with k-Nearest Neighbors/Naive Bayes and Normalized HMMs with k-Nearest Neighbors) we decided to also report measurements about the area under the Precision-Recall curve, which can be found in table 2.

In order to be consistent with related works despite using 3-mer models in almost the entirety of this work, we provide a comparison between the Normalized Bag-of-words models using k-mers of the size of 2 and 3 respectively.

These results (shown in Table 3 indicate that there isn’t much difference between the two models in terms of the metrics we used, therefore leading us to believe that the results provided in our work are in accordance with those of previous works that relied heavily on 2-mer models.

Lastly, in order to evaluate the validity of the above analysis of our experimental results we created a dataset which consisted of tuples of the form (*Representation, Algorithm, Fold*#, *Metric*) for each of the three aforementioned metrics that we used and, by applying the ANOVA (Analysis of Variance), we measured the effect of each combination of the above first three independent variables on the performance of our model in respect to the relative metric, which acted as the fourth and dependent variable.

Starting from the results of the ANOVA test (Table 4), we found that the main effects of both the types of algorithms and representations used were statistically significant in the performance of our model, while the effect of the fold number (*F old*#) was statistically insignificant, therefore not affecting the performance of our experiments. For this specific test we assumed a level of significance equal to 0.05, thus rejecting the null hypothesis *H*_0_, should the respective p-value be less than 0.05. Finally, the interaction effect between different representations and classifiers was, as expected, statistically significant.

Furthermore, in order to single-out the representation that achieved the best performance, we ran Tukey HSD (Honest Significant Difference) tests on the aforementioned dataset we created (Table 5). This analysis aims to show which configuration is statistically significantly better than others. Each formed group — identified by a lowercase letter — indicates a set of configurations with (statistically) equivalent performance.

**Table 5:** Results of Tukey test on the performance of different representations with respect to the three different metrics used. Representations are ordered alpharithmetically in terms of performance (i.e., b is considered better than a). Measurements which are within statistical significance to the best are underlined while the best measure overall appears in bold.

| Metric | Representation | Group |
| --- | --- | --- |
| Recall | Centroid Bag-of-words | **c** |
|  | HMM | a |
|  | N-gram graphs | **c** |
|  | Normalized Bag-of-words | <u>bc</u> |
|  | Normalized HMM | b |
| Precision | Centroid Bag-of-words | c |
|  | HMM | a |
|  | N-gram graphs | **d** |
|  | Normalized Bag-of-words | <u>cd</u> |
|  | Normalized HMM | b |
| F-measure | Centroid Bag-of-words | c |
|  | HMM | a |
|  | N-gram graphs | **d** |
|  | Normalized Bag-of-words | c |
|  | Normalized HMM | b |

Our results failed to determine a single, prevalent classification algorithm in terms of performance, although the Decision Trees (*DT*) and Support Vector Machines (*SV M*) seemed to stand out in some cases (Table 6). Last but not least, we evaluated the performance of the Representation/Classification algorithm pairs used (Tables 7-9) and, ultimately, saw that the N-gram graphs tend to be successful regardless of the classification algorithm applied, whereas Baseline Bag-of-Words tend to find success only when used in conjunction with certain classifiers. Other representations which also challenge the aforementioned two are, in certain limited situations, Bag-of-Words with reduced dimensions and Normalized Hidden Markov Models. Thus, overall N-gram graphs demonstrate high robustness with consistently high performance.

**Table 6:**
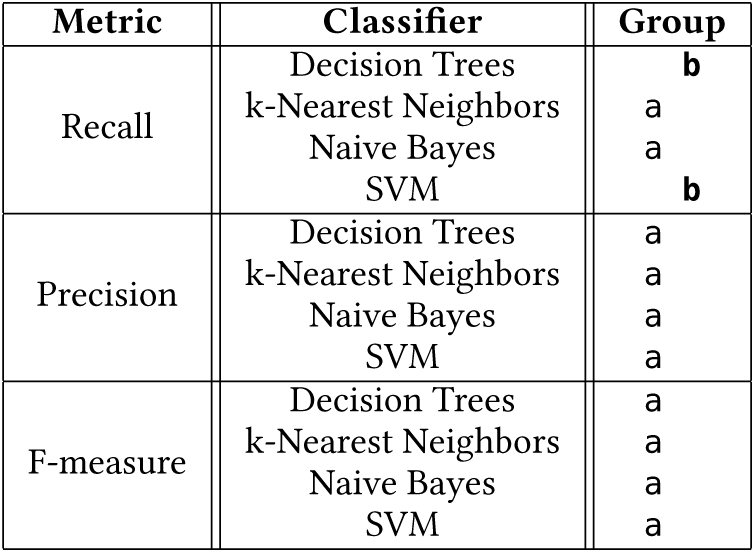
Results of Tukey test on the performance of different classification algorithms with respect to the three different metrics used. Classification algorithms are ordered alpharithmetically in terms of performance (i.e., b is considered better than a). Measurements which are within statistical significance to the best are underlined while the best measure overall appears in bold.

| Metric | Classifier | Group |
| --- | --- | --- |
| Recall | Decision Trees | **b** |
|  | k-Nearest Neighbors | a |
|  | Naive Bayes | a |
|  | SVM | **b** |
| Precision | Decision Trees | a |
|  | k-Nearest Neighbors | a |
|  | Naive Bayes | a |
|  | SVM | a |
| F-measure | Decision Trees | a |
|  | k-Nearest Neighbors | a |
|  | Naive Bayes | a |
|  | SVM | a |

**Table 7:** Results of Tukey test on different Representations / Classification algorithms with respect to the Recall metric. Representation/Classification Algorithm pairs are ordered alpharithmetically in terms of performance (i.e., b is considered better than a). Measurements which are within statistical significance to the best are underlined while the best measure overall appears in bold.

| Representation | Classification Algorithm | Group |
| --- | --- | --- |
| Centroid Bag-of-words | Decision Trees | <u>fg</u> |
|  | k-Nearest Neighbors | ce |
|  | Naive Bayes | <u>eg</u> |
|  | SVM | **g** |
| HMM | Decision Trees | **g** |
|  | k-Nearest Neighbors | b |
|  | Naive Bayes | a |
|  | SVM | **g** |
| N-gram graphs | Decision Trees | <u>fg</u> |
|  | k-Nearest Neighbors | <u>eg</u> |
|  | Naive Bayes | <u>fg</u> |
|  | SVM | **g** |
| Normalized Bag-of-words | Decision Trees | c |
|  | k-Nearest Neighbors | def |
|  | Naive Bayes | **g** |
|  | SVM | **g** |
| Normalized HMM | Decision Trees | <u>eg</u> |
|  | k-Nearest Neighbors | b |
|  | Naive Bayes | cd |
|  | SVM | **g** |

**Table 8:** Results of Tukey test on different Representations / Classification algorithms with respect to the Precision metric. Representation/Classification Algorithm pairs are ordered alpharithmetically in terms of performance (i.e., b is considered better than a). Measurements which are within statistical significance to the best are underlined while the best measure overall appears in bold.

| Representation | Classifier | Group |
| --- | --- | --- |
| Centroid Bag-of-words | Decision Trees | g |
|  | k-Nearest Neighbors | <u>gh</u> |
|  | Naive Bayes | f |
|  | SVM | f |
| HMM | Decision Trees | a |
|  | k-Nearest Neighbors | c |
|  | Naive Bayes | de |
|  | SVM | a |
| N-gram graphs | Decision Trees | <u>gh</u> |
|  | k-Nearest Neighbors | **h** |
|  | Naive Bayes | <u>gh</u> |
|  | SVM | <u>gh</u> |
| Normalized Bag-of-words | Decision Trees | f |
|  | k-Nearest Neighbors | <u>gh</u> |
|  | Naive Bayes | f |
|  | SVM | **h** |
| Normalized HMM | Decision Trees | d |
|  | k-Nearest Neighbors | e |
|  | Naive Bayes | de |
|  | SVM | b |

In the Discussion section, we also comment on the fact that representative-based approaches can have — within statistical significance — equivalent performance to high dimensionality, instance-based representations. However, this does not happen regardless of the classification algorithm in most cases (cf. Table 9).

**Table 9:** Results of Tukey test on different Representations / Classification algorithms with respect to the F-measure metric. Representation/Classification Algorithm pairs are ordered alpharithmetically in terms of performance (i.e., b is considered better than a). Measurements which are within statistical significance to the best are underlined while the best measure overall appears in bold.

| Representation | Classifier | Group |
| --- | --- | --- |
| Centroid Bag-of-words | Decision Trees | hi j |
|  | k-Nearest Neighbors | ghi |
|  | Naive Bayes | fh |
|  | SVM | fg |
| HMM | Decision Trees | a |
|  | k-Nearest Neighbors | bc |
|  | Naive Bayes | b |
|  | SVM | a |
| N-gram graphs | Decision Trees | <u>jk</u> |
|  | k-Nearest Neighbors | <u>ik</u> |
|  | Naive Bayes | <u>ik</u> |
|  | SVM | <u>ik</u> |
| Normalized Bag-of-words | Decision Trees | f |
|  | k-Nearest Neighbors | hi j |
|  | Naive Bayes | gh |
|  | SVM | **k** |
| Normalized HMM | Decision Trees | e |
|  | k-Nearest Neighbors | d |
|  | Naive Bayes | e |
|  | SVM | c |

**Table 10:** 2-mer and 3-mer sorted by their average rank, with respect to their mean information gain across a 10-fold cross-validation evaluation.

| # | 2-mers |  |  | 3-mers |  |  |
| --- | --- | --- | --- | --- | --- | --- |
|  | name | rank | inf. gain | name | rank | inf. gain |
| 1 | TT | 0.20 | 0.55 | CCA | 0.20 | 0.33 |
| 2 | TG | 0.80 | 0.55 | GGC | 0.80 | 0.32 |
| 3 | GG | 2.00 | 0.54 | AGC | 2.70 | 0.30 |
| 4 | AT | 3.20 | 0.52 | TAA | 3.00 | 0.30 |
| 5 | GC | 4.40 | 0.52 | AAC | 4.90 | 0.29 |
| 6 | AC | 4.60 | 0.52 | CAA | 5.00 | 0.29 |
| 7 | AG | 6.20 | 0.51 | CTA | 5.20 | 0.29 |
| 8 | CG | 6.90 | 0.51 | TTA | 7.80 | 0.29 |
| 9 | CT | 7.70 | 0.51 | ATT | 8.70 | 0.28 |
| 10 | TC | 9.10 | 0.48 | AAA | 9.60 | 0.28 |

### 3.3 Discriminative features and patterns

In order to illustrate discriminative n-mers in the input sequences, we performed a number of additional evaluations on the Normalized BoW representation approach. This feature encoding method was used because it is the only method where n-mer information from the input sequence is directly encoded in the representation vector, rather than in a similarity to the representative, as is the case for the other approaches. To this end, we measured the mean information gain for each 2-mer and 3-mer in the input data under a 10-fold cross-validation scheme. In table 3.2 we list the 10 most important 2-mers and 3-mers, ranked with respect to the mean rank across all folds (column *rank*), and with the mean information gain value (column *inf. gain*).

For both k=2 and k=3, k-mers containing the TT and GG dinucleotides are showing up among the ones with the greatest information gain, while TA containing triplets are also enriched in the top of the list. TT/AA and GG/CC are known to carry particular structural properties that confer rigidity (TT/AA) and flexibility (GG/CC) to the double helix. Their counter-phase periodical distribution has been suggested to be one of the main determinants of nucleosome positioning sequences. On the other hand, the TA dinucleotide is the one exhibiting a characteristic structural propensity, called “propeller twist” which is expected to stabilize the DNA helix in a rigid conformation. Thus it appears that our analysis may reveal different structural constraints of the DNA reflected on the genomic sequences under study.

## 4 Discussion

The results of our analyses may be summarized in the following points: First, that there are indeed content constraints in genomic sequences that may be used as determinants of nucleosome positioning and that these may be extended beyond the well-reported homodimer periodicities (AA/TT) or an overall preference for high GC%. Second, that such sequence determinants are not entirely reflected in k-mer occurrences but also on the relative positions thereof. This is supported by the superior perfomance of the N-gram Graphs representations, when compared to both k-mer-only approaches (Bag-of-words) and probabilistic approaches implying nearest-neighbour dependencies (HMMs). The results presented herein, suggest that the underlying sequence constraints are at the same time subtle and long-ranged and thus better captured by an approach like N-gram Graphs. Last but not least, through extensive combinations of different representations with various classification methods we show that the choice of both is not trivial and that different classification approaches may be more suitable depending on the representation used.

Our results indicated that the N-gram graphs and Bag-of-words models are better suited for our problem than HMM. This is better shown in **section 5.2** where comparisons between the different models are made, based on the Recall, Precision and F-measure metrics. The prevalence of these models indicates that the exact order of nucleotides or n-mers in the genomic sequence does not appear to play a central role in controlling the formation of nucleosomes, a feature which is rather attributed to the presence (or absence) of certain nucleotides or n-mers. This is consistent with the general notion of nucleosome-forming sequences being mostly governed by simple compositional rules such as dinucleotide periodicities and over/under-representations of k-mers like the ones we report here. This apparent lack of an internal “structure” is at the same time the basis of the assumed “nucleosomal code’s” flexibility and of the inherent difficulty in nucleosome positioning determination from the underlying sequence. Normalized HMMs, on the other hand, tend to achieve better performance than their plain counterparts, therefore strengthening our intuition that certain n-mers of limited length tend to correlate with nucleosome positioning, as in this case the predictive strength of a sequence diminishes as it grows in length.

A fact which is worth mentioning is that our dataset exhibited class imbalance, having 1033 instances for the NFR class and 2660 instances for the NOS class. This 1:2 ratio of NFR to NOS instances indicates that the random chance performance of any model trained on our dataset would be around 72%, thus enabling us to note that the performance of the model trained on the plain HMM representation features performance that is marginally better than that. This leads us to conclude that the plain HMMs suffer from the Most Common Class problem, whereas all the other models, the Normalized HMM included, fared better against these problems and proved to tackle them efficiently.

At the same time, the success of the N-gram Graphs representation clearly suggests that a — limited in extent — internal structure does exist in nucleosomal sequences. What the performance of N-gram Graphs shows is that certain “neighborhoods” of n-mers can identify NFR or NOS genomic sequences, therefore suggesting that relative n-mer positions may be informative of the sequence’s affinity for the histone octamer. In the past, the representation of DNA sequences with the use of N-gram Graphs has only been used once in the case of conserved non-coding sequences with promising results [27]. In this work, the potential of N-gram Graphs to capture subtle sequence constraints related to complex functional roles is further demonstrated.

What is noteworthy concerning the representative-based approaches is that they drastically reduce the feature space, while retaining equivalent level of performance (within statistical significance) to their instance-based counterparts. On the other hand, the preprocessing in the representative-based approaches is significantly increased (since we need to extract the representatives), which reduces the overall speed gain.

All in all, as it was experimentally shown in the Evaluation section both regarding the performance metrics and the ANOVA/Tukey tests, there exists an apparent difference in the efficiency with which different representation/classifier pairs tackle the problem of predicting nucleosome positions in the genome. These results further support our intuition that the selection of the representation/classifier pair singificantly affects performance. As a matter of fact, the selection of the representation in which our problem was to be modeled was more significant than the selection of the classification algorithm (cf. Table 4). This leads us to the conclusion that monitoring the selection of these variables, particularly of the representation and secondarily of the classification algorithm, should not be neglected in any similar studies in the future, as it appears to play a vital role in the resulting performance.

## 5 Concluding Remarks

In this study our aims were twofold. On one hand we sought to elaborate and extend findings of previous works on the existence of particular sequence prerequisites guiding nucleosome positioning. On the other hand we set out to evaluate a variety of Machine Learning algorithms and representations in the definition of such sequence determinants and the extraction of contextual information. Our analyses provide us with a clearer intuition about the presence of certain rules guiding the positioning of nucleosomes. Moreover, the success of certain models in this selection sheds a new light on certain properties of the DNA sequence that have a greater impact in regulating these positions. A distinguishing feature of our work is the demonstration of N-gram Graphs as a preferable representation of genomic sequences. N-gram Graphs are, by construction, able to capture both compositional over-/under-representations and positional constraints between the various n-grams in a way that is more flexible than usual probabilistic approaches such as HMMs. The N-gram Graphs superior perfomance against HMMs, in the context of nucleosome positioning sequences is a clear indication of their potential in the study of genomic sequences.

In the ongoing race of accumulating data and method development, our study brings an additional aspect into the spotlight. That is the importance of the combination of different representations with different classification methods. As results shown in tables 6 to 8 suggest, the use of different classification methods appear to be favorable to different representations and thus, future works on sequence analysis employing machine learning analysis may need to consider this additional level of complexity.

Last but not least, our work points out that the development of novel representations of sequence content for a wider variety of possible features is likely to further optimize the feature extraction process. Better and more carefully conceived features can ultimately lead to better classification results, which can capture more accurately the factors that distinguish the functionality of different genomic sequences.

In the future, we will need to devise a way to identify representative features from representative-based representations. This will allow better explainability of the classificiation results, while retaining an efficient vector space.

## Acknowledgements

**Funding**

(Acknowledgements for funding sources will be provided in the camera-ready version, if needed).

## Footnotes

3 https://github.com/aubry74/Jahmm

4 https://sourceforge.net/p/jinsect/wiki/Home/

5 http://www.cs.waikato.ac.nz/ml/weka/

